# Screening de. novo designed protein binders in unpurified lysate using flow induced dispersion analysis

**DOI:** 10.1101/2025.06.17.660127

**Authors:** Francisca Pinheiro, Jan S. Nowak, Elena Zueva, Emily C. Pheasant, Ida Kjærsgaard Grene, Vili Lampinen, Magnus Kjaergaard

**Author notes:** For correspondence: Magnus Kjaergaard.

## Abstract

Computational protein design can create binders against targets of interest, but identifying binders with sufficient affinity still requires biochemical screening of many designs. In this work, we test flow-induced dispersion analysis (FIDA) as a method for screening binders in a time and cost-effective manner. FIDA uses Taylor dispersion analysis to determine the hydrodynamic radius of fluorescently labelled biomolecules and their complexes. Here, we use FIDA to assess the binding of RFdiffusion-designed protein binders against the small helical peptide ALFA-tag and the GK domain of PSD-95. Successful binders can be identified in a single measurement using heat-treated bacterial lysates allowing rapid identification of binders with high affinity, and thermostability. Subsequent titration experiments show micromolar affinities for ALFA-tag binders and nanomolar affinities for GK domain binders. The lack of immobilization, the minimal sample volume, and the compatibility with complex biological samples, positions FIDA as a valuable tool for the screening and characterization of computationally generated protein binders.

## Introduction

Protein design has been revolutionized by deep learning algorithms^1,2^. There are various algorithms which can generate novel protein folds, tailored to specific functions, and design sequences that are predicted to adopt the structure of these folds. Together, these algorithms enable the design of plausible, functional proteins in.silico. One of the most common goals with in.silico design is to generate synthetic protein binders that bind strongly and specifically to a desired target. This has become possible using deep learning models, such as RFdiffusion^3^ or BindCraft^4^. In addition to their binding abilities, de.novo designed proteins often have desirable secondary properties such as high thermostability and expression yield in.E¡.coli^3^.

Experimental validation and characterization are necessary parts of protein design campaigns but have become their main bottleneck. For binder design using RFdiffusion and proteinMPNN, an average of 19 % of binder candidates show binding at 10 µM across 5 targets^3^. However, far fewer have sufficient affinity for most common applications and, as a result, binder design projects still require the screening of tens to hundreds of candidates. There is thus a need for rapid and cost-effective methods for screening large numbers of binder designs.

An ideal screening method for binders should require minimal sample amounts and little to no sample pre-treatment. Bacterial expression systems are particularly convenient, as bacteria can be grown in 24- or 96-well plates using a culture volume of approximately two millilitres per well. Depending on the downstream screening method used, various degrees of purification can be performed – e.g. nickel-affinity chromatography in a 96-well format and subsequent size-exclusion chromatography using an automated sample loader. However, these steps require considerable labour and infrastructure, and it is desirable to avoid them during initial screening stages. Screening for binding is typically done by a single-point biolayer interferometry (BLI)^3^, which measures protein adhesion to a target immobilized on a hydrogel matrix. BLI is favoured over other techniques due to its ability to measure multiple samples in parallel and its tolerance for crudely purified samples. However, it has some drawbacks such as a high false positive rate due to nonspecific interactions with the matrix and the requirement for immobilisation of the target protein. Additionally, BLI does not provide any further information about the complex formed.

In this work, we explore a newly developed technique called flow induced dispersion analysis (FIDA) for the screening of de.novo designed protein binders. FIDA is a microfluidic technique used for size-based characterization of protein (un)folding, stability and binding^5^. Briefly, FIDA employs Taylor dispersion analysis to determine the diffusion coefficients of molecules, which are then used to calculate the apparent hydrodynamic radius (R_h_) via the Stokes-Einstein equation^6^. In practice, a sample is injected under pressure into a fused silica capillary (typically 75 µm in diameter), and the fluorescence intensity of a labelled molecule within the sample - referred to as the indicator - is measured at the end of the capillary, producing a symmetrical, bell-shaped peak (**Fig. 1A**). The resulting curve can be fitted with a Gaussian function to extract the diffusion coefficient, and by extension, the R_h_. Upon binding, the diffusion rate of the fluorescent molecule typically decreases, resulting in a broader peak, and its R_h_ increases (**Fig. 1B**). Experimentally obtained R_h_ values can be compared to the expected R_h_, which can be calculated from the predicted structure of the complex^7^. Experiments can be performed in the presence of increasing analyte concentrations to determine the dissociation constant (K_D_) or as a single-point measurement for screening.

**Fig. 1.**
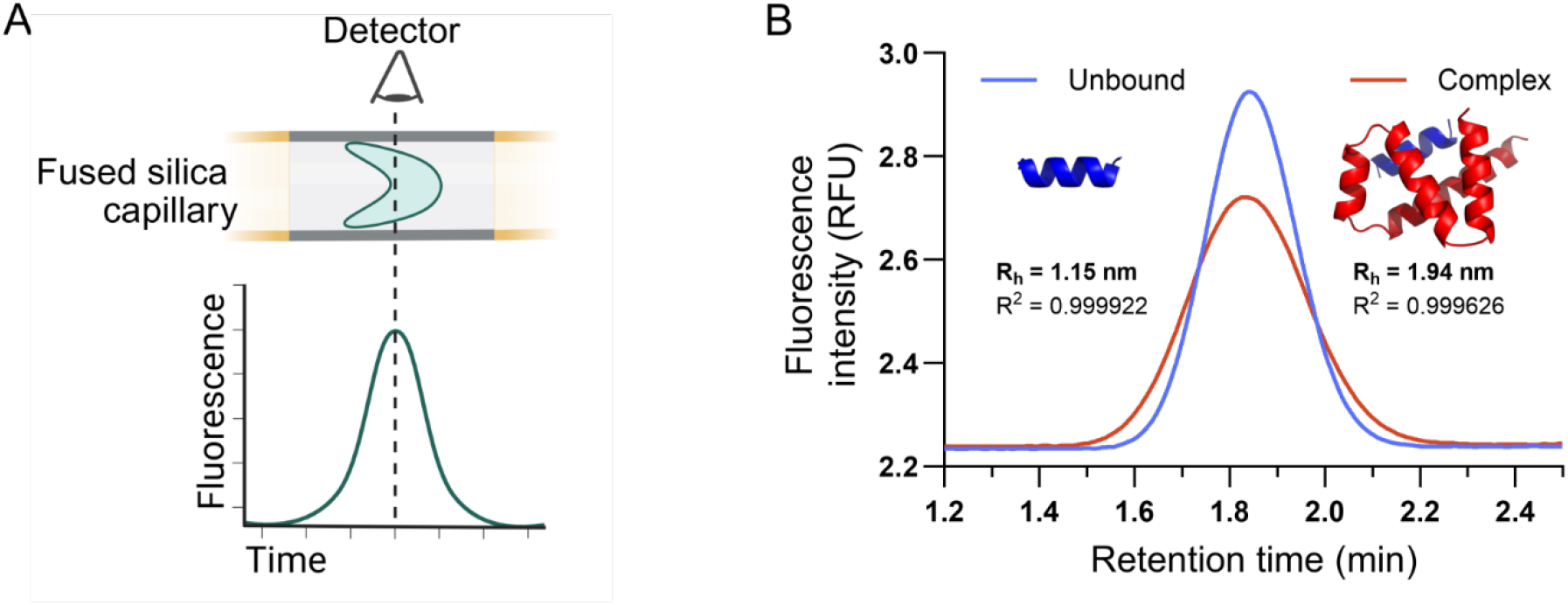
Schematic of the FIDA method for detecting binding. (A) In FIDA, a bell-shaped peak, or Taylorgram, is produced when the fluorescently labelled molecule (indicator) reaches the detector. (B) Taylorgrams for unbound ALFA-tag indicator (in blue) and in complex with an analyte (10 µM binder 5A in red), show how the width of the peak changes upon binding.

Here, we have designed binders against a 2 kDa peptide and a 34 kDa globular domain and tested FIDA’s ability to efficiently screen for complex formation. We show that binders can be identified in a single measurement, using heat-treated bacterial lysates, effectively eliminating the need for protein purification. Consequently, primary hits can in principle be identified within one day of binder expression, significantly accelerating the screening process. FIDA’s compatibility with complex samples, its low sample consumption and minimal need for optimization, positions it as a promising platform for screening de.novo designed binders.

## Methods

### Preparation of DNA constructs

The DNA constructs coding for ALFA-tag and PSD-95 GK-domain binders were obtained from Twist Bioscience (South San Francisco, CA, USA). The coding regions were cloned into pET-28a(+) vectors using the NdeI/XhoI restriction sites. As a result, all protein binders contained a 6xHis-tag in the N-terminus, which was separated from the binder sequence by a thrombin site. The DNA construct encoding the SH3-GK module of PSD-95 was obtained from Genscript (Piscataway, NJ, USA). Residues 431-724 of human PSD-95 (Uniprot: P78352) were synthesised with an N-terminal TEV site and cloned into the NdeI/BamHI restriction sites of pET15b, generating a construct with a TEV-cleavable 6xHis-tag. Protein sequences are detailed in the supplementary material.

### Expression and purification of protein binders

All protein binders were expressed in Escherichia. coli BL21(DE3) cells in 50 mL ZYM-5052^8^ supplemented with 50 µg/mL kanamycin at 37°C for 20 – 22 h. The cells were harvested by centrifugation and lysed either by chemical (ALFA-tag binders) or heat (PSD-95 GK-domain binders) lysis. For chemical lysis, the bacterial pellets were resuspended with 2.5 mL B-PER™ reagent (Thermo Fisher Scientific, Waltham, MA, USA) supplemented with 0.2 mM PMSF (Carl ROTH, Karlsruhe, Germany) and 10 µg/mL DNAseI (Roche, Basel, Switzerland), and incubated for 15 min at room temperature. For heat lysis, the bacterial pellets were resuspended in 20 mM sodium phosphate buffer pH 7.6, 300 mM NaCl, 0.2 mM PMSF, and incubated for 15 min at 95 °C. In both cases, the soluble fraction was recovered by centrifuging the lysates at 3,041 x.g for 20 min and applied to gravity flow columns packed with Ni Sepharose 6 Fast Flow (Cytiva, Uppsala, Sweden), which were previously equilibrated with 20 mM sodium phosphate pH 7.6, 300 mM NaCl, 5 mM imidazole. The columns were washed with buffers (20 mM sodium phosphate buffer pH 7.6, 300 mM NaCl) containing 20 mM imidazole and eluted with 300 mM imidazole. The elution fractions were dialysed against 20 mM sodium phosphate pH 7.6, 300 mM NaCl and were either subjected to thrombin cleavage (ALFA-tag binders) or directly concentrated using Amicon® Ultra 15 mL Centrifugal Filters (Merck Millipore, Burlington, MA, USA) with 3 kDa MWCO (PSD-95 GK-domain binders).

The N-terminal His-tag of ALFA-tag binders was removed by incubating the purified binders with thrombin (2 U/mg of protein) overnight at room temperature and applying them to gravity flow columns packed with nickel sepharose as before. As expected, SDS-PAGE gels showed purified binders in the fractions containing 5 and 20 mM imidazole, which were then combined and dialysed against 20 mM sodium phosphate buffer pH 7.6, 300 mM NaCl. Finally, the proteins were concentrated as described for the PSD-95 GK-domain binders.

After concentration, both ALFA-tag and PSD-95 GK-domain binders were flash frozen in liquid nitrogen and stored at –20 °C until used. Protein concentration was determined spectrophotometrically at 280 nm, or 205 nm, for binders containing no tryptophan or tyrosine as described in Anthis and Clore (2013)^9^.

### Expression and purification of the PSD95 SH3-GK module

The SH3-GK module of PSD-95 was expressed in E¡. coli BL21(DE3) cells in ZYM-5052 autoinduction medium supplemented with 100 µg/mL ampicillin. Expression was carried out for 42 h, of which the first 3 h were at 37 °C, after which the temperature was lowered to 20 °C. The cells were then harvested by centrifugation (6,000 ≥.g, 20 min, 20 °C) and lysed by sonication. Briefly, cell pellets were resuspended in 20 mM sodium phosphate, 500 mM NaCl, 0.1 mM TCEP, pH 7.4, supplemented with PMSF (0.2 mM) and pepstatin A, chymostatin and leupeptin (1 μg/mL each) and subsequently sonicated while on ice. For purification, the soluble fraction was separated by centrifugation (22,789 ≥.g, 15 min, 4 °C) before being applied to a gravity-flow column packed with Ni-Sepharose 6 Fast Flow (Cytiva) which was previously equilibrated with sodium phosphate buffer containing 5 mM imidazole (20 mM sodium phosphate, 500 mM NaCl, 5 mM imidazole, 0.1 mM TCEP, pH 7.4). The column was then washed with buffer containing 20 mM imidazole and His-tagged SH3-GK was eluted by increasing imidazole concentration to 500 mM.

To remove the His-tag, the elution fraction was spiked with TEV protease (1:100 m/m) and dialysed (overnight, 4 °C) against 20 mM sodium phosphate, 500 mM NaCl, 1 mM TCEP, pH 7.4. Cleaved SH3-GK was isolated by applying the TEV-treated and dialysed sample to a gravity-flow column packed with Ni-Sepharose as before and collecting the flow-through and the wash. Finally, SH3-GK without His-tag was concentrated using a Vivaspin® 20 centrifugal concentrator (Sartorius AG, Göttingen, Germany) with a 10 kDa MWCO, and purified by size-exclusion chromatography on a Superdex 75 Increase column (Cytiva) in 20 mM sodium phosphate, 500 mM NaCl, 0.1 mM TCEP, pH 7.4, at room temperature. Chromatography fractions were analysed by SDS-PAGE and the purest fractions were flash-frozen and stored at -70 °C.

### Preparation of bacterial lysates

For comparison of lysis methods, bacterial cultures expressing binders were split in half and subjected to chemical or heat lysis as previously explained, and the samples were centrifuged at 3,041 x.g for 20 min. The supernatants (soluble fraction) were aliquoted, flash frozen in liquid nitrogen and stored at –70 °C until used for FIDA experiments.

### FIDA experiments

All experiments were conducted on a FIDA 1 instrument (Fida Biosystems ApS, Copenhagen, Denmark) using light-emitting diode (LED) induced fluorescence detection with an extinction wavelength of 480 nm and an emission wavelength of >515 nm. Permanently coated capillaries were used throughout this study to reduce stickiness (75 μm inner diameter, 375 μm outer diameter, 1 m total length, and 84 cm length to detection window). The assay buffer was 20 mM sodium phosphate buffer pH 7.6, 300 mM NaCl + 0.05% pluronic acid F-127 (Invitrogen, Thermo Fisher Scientific). All measurements were done using the capillary mixing approach and in triplicate. The expected R_h_ was calculated from the structure of the complexes, which was predicted with either AlphaFold2 (ALFA-tag) or AlphaFold3 (SH3-GK)^10^, using the PDB correlator that is available in the FIDA software v3.1^7^.

#### Indicator preparation

N-terminally FITC-labelled ALFA-tag (SRLEEELRRRLTE) was ordered from GenScript at 95% purity. Lyophilized pure peptide was resuspended in 50% DMSO to a concentration of 4 mM, flash frozen with liquid nitrogen and stored at -70°C until assayed. PSD-95 SH3-GK was mixed with Alexa Fluor 488 (AF488) NHS Ester (Invitrogen, Thermo Fisher Scientific) at a molar protein:dye ratio of 1:3.4 and with sodium bicarbonate at a final concentration of 100 mM. After 1 h incubation at room temperature, excess free, unreacted dye was removed using a CentriPure 25 ‒ Z25M column (emp BIOTECH GmbH, Berlin, Germany). The concentration of labelled protein was measured using the Labelled Proteins application in a DS-11 Spectrophotomer (DeNovix, Wilmington, DE, USA), assuming an extinction coefficient of 42400 M_-1_ cm_-1_ for the SH3-GK protein. AF488-labelled SH3-GK was flash frozen in liquid nitrogen and stored at -70°C until used.

#### Assay sequence

All FIDA experiments were done using the standard coated capillary method, which consists of four automated steps necessary to acquire a single measurement: 1) a 75 s pre-equilibration at 3500 mbar with the running buffer, 2) 20 s analyte injection at 3500 mbar, 3) 10 s injection of the indicator with or without analyte at 50 mbar, and 4) 180 s subsequent injection of the analyte at 400 mbar. Data recording was done in the last step.

#### Binding screening and affinity determinations

Binding screening against ALFA-tag was performed in a capillary mixing (capmix) mode by loading 10 µM purified binders in steps 2 and 4 of the sequence and 50 nM FITC-ALFA in step 3. The R_h_ values were calculated by processing the Taylorgrams using the FIDA Software v3.1 (Fida Biosystems) with a Taylorgram fraction of 75%. The affinity of the purified binders to their corresponding targets was also assessed using capmix mode, by loading different concentrations of the binders in steps 2 and 4 and injecting FITC-ALFA (50 nM) or AF488-labelled SH3-GK (80 nM) in step 3. The R_h_ values were determined as before, except that for SH3-GK the Taylorgrams were fitted for multiple species with the R_h_ of the second species locked to 0.6 nm to account for free label. Determination of the dissociation constants (K_D_) was done by fitting the titration curves to the 1:1 binding stoichiometry model that is available on the FIDA software (V3.1)^11^.

#### Screening on bacterial lysates

Binding screening using chemical (only for ALFA-tag) and heat lysates was done in capmix mode by injecting the lysates in both loading steps and 200 nM FITC-ALFA/AF488-labelled SH3-GK as indicators. The lysates were diluted to different extents using assay buffer before being injected. In both cases, a sample of E¡.coli.that does not express any binder was lysed in the same manner and used as control. The size of FITC-ALFA and AF488-labelled SH3-GK in the absence or presence of binders was determined as explained above. The results obtained for the binding screening in heat lysates from bacteria expressing the PSD-95 GK binders were analyzed by one-way ANOVA Dunnett’s multiple comparisons test using GraphPad Prism 10.5.0. p < 0.05 was considered statistically significant (* statistically significant at p < 0.05; ** statistically significant at p < 0.01; *** statistically significant at p < 0.001).

## Results

### De.novo design of binders against the ALFA-tag

As a test case for the binder screening, we used the ALFA-tag as target epitope. The ALFA-tag is a designed epitope-tag that forms an amphipathic helix and has favourable properties for imaging such as the absence of overall charge and lysine residues^12^. It is an idealized test case of the short linear motifs frequently found in intrinsically disordered regions. The ALFA-tag is recognized by a nanobody with picomolar affinity^12^ through interactions with its hydrophobic face (**Fig. 2A**).

**Fig. 2.**
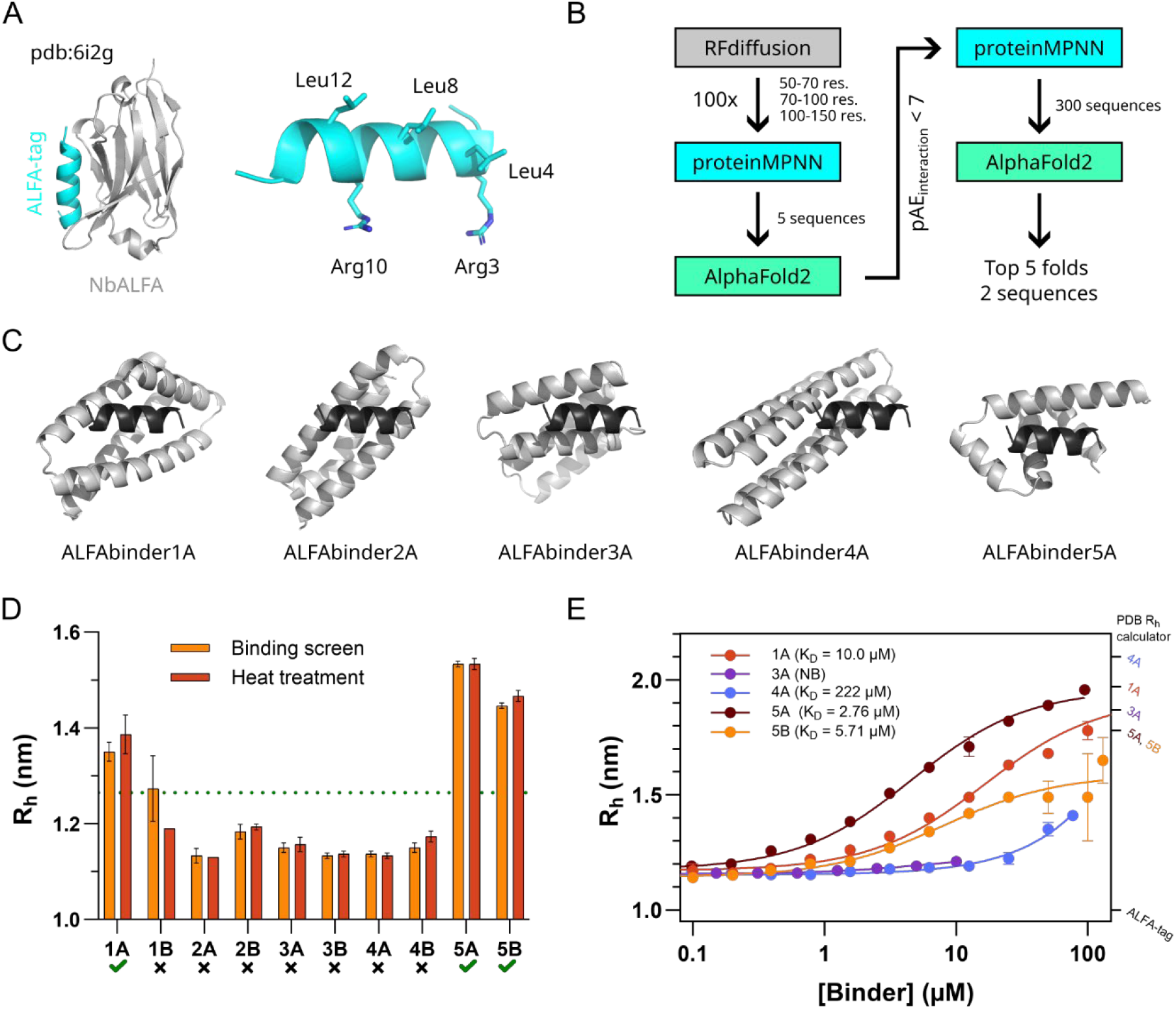
Design and characterization of ALFA-tag binders. (A) Crystal structure of ALFA-tag (in cyan) in complex with the nanobody (in grey) that was used as a starting point for binder design. The ALFA-tag alone is shown on the right with the epitope residues depicted as sticks. (B) Schematic overview of the de.novo protein binder design pipeline. (C) Complexes of the ALFA-tag (in black) with the designed binders (in grey) as predicted by AlphaFold2. For each fold, only one of the ALFA-tag:binder complexes is shown as their structure is identical. (D) Apparent R_h_ of FITC-ALFA (50 nM) when mixed with 10 µM purified binder inside the capillary (light orange). To assess the thermal stability of the binders, the binders were heated at 95°C for 5 min and FIDA measurements were performed in the same way (dark orange). An increase in R_h_ of 10% relative to the R_h_ obtained for FITC-ALFA alone was selected as the cutoff for binding and is indicated with a green dotted line. The green ticks highlight the positive hits. (E) Binding curves generated by representing the apparent R_h_ of FITC-ALFA as a function of binder concentration. The solid lines correspond to the fitting to the 1:1 binding isotherm. The predicted sizes for ALFA-tag alone or in complex with the different binders are indicated in the right y-axis. All R_h_ values correspond to mean ± SD (n = 3).

We extracted the coordinates of the ALFA-tag from the PDB file of the ALFA-tag:nanobody complex^12^ and used them as a template for RFdiffusion/proteinMPNN^3,13^ binder design (**Fig. 2B**). The target epitope consisted of the three hydrophobic residues (L4, L8 and L12), supplemented with arginine residues from one side of the helix (R3, R10) to induce a larger interface. Designs were computationally scored by complex prediction by AlphaFold2^14^ resulting in an in.silico success rate of about 5% with a threshold of pAE_interaction_ < 10. Folds with pAE_interaction_ below 7 were inspected and folds with only two helices were removed due to concerns about stability and interface size. The remaining folds were resampled 300 times in proteinMPNN, and for each of the top 5 folds, the two sequences with the lowest pAE_interaction_ were selected for characterization (**Fig. 2C**).

Codon-optimized DNA sequences of the 10 binder candidates were synthesized with an N-terminal thrombin cleavable His-tag and expressed in E¡. coli in autoinduction media^8^. The resulting proteins were purified by Ni-NTA affinity chromatography with a 1-step elution. All 10 binders resulted in soluble expression and proteins that could be purified although with varied yield (3-148 mg protein per litre of culture).

### Screening and titration of purified ALFA-tag binders by FIDA

To assess the binding of de.novo designed proteins to ALFA-tag, we used a N-terminal FITC-labelled ALFA peptide as the observable indicator which allowed us to follow the fluorescence at 480 nm. 50 nM FITC-ALFA resulted in a robust signal (S/N >50) in dilute buffers and a R_h_ of 1.15 nm, which is close to the predicted value of 1 nm based on the PDB structure without the FITC modification. Next, we performed the experiment in the presence of the 10 designed binders at a concentration of 10 µM. The experiments were conducted in a capillary mixing (capmix) mode, where the proteins are mixed inside the capillary. Capmix experiments are faster to set up and require smaller amounts of indicator, but it is not guaranteed that slow associating interactions have reached equilibrium at the end of the capillary (reaction time < 1.8 min). This means that binders with slow association kinetics may not be detected. The binders that induced an increase in R_h_ equal or greater than 10% relative to the previously measured for FITC-ALFA alone (1.15 nm) were considered positive hits (**Fig. 2D**). Binders 1A, 1B, 5A and 5B fulfilled this condition, although the error associated with 1B was relatively high.

Designed proteins are often highly thermostable such that many do not denature at temperatures below the boiling point of water^3^. In many applications, thermostability is a desirable secondary trait for a designed binder, and it is thus useful to be able to rapidly screen for stability. We screened for thermostability by incubating the binders for 5 minutes at 95 °C before the FIDA measurements. Retained binding indicates that the protein either does not denature at 95 °C or refolds without aggregating. When this analysis was applied to the ALFA-tag binders, we observed that 1A, 5A, and 5B retained their ability to increase the measured R_h_ of the ALFA-tag (**Fig. 2D**), while a much lower R_h_ was observed for 1B. This indicates that this binder is susceptible to heating or that it was a false positive in the initial screen, which is further supported by the large error associated with the R_h_ calculation. In either case, this strategy allowed us to confirm that three of the potential binders are positive hits while also being thermostable.

To quantify the affinity of the binders, we titrated the three positive hits (1A, 5A, and 5B) and two non-hits (3A and 4A) by a two-fold serial dilution of the binders ranging from the highest available concentration (typically ∼100 µM) down to 0.1 µM (**Fig. 2E**). Titration of all three positive hits produced classical isotherms that could be fitted to a 1:1 binding model^11^, resulting in K_D_ values of 10.0 ± 1.46 µM, 2.76 ± 0.25 µM, and 5.71 ± 1.50 µM, for 1A, 5A, and 5B, respectively. As expected, the negative controls resulted in either high or no measurable K_D_. The sigmoid binding curves indicate that the interactions identified by FIDA analysis are specific and saturable, ruling out non-specific interactions.

A potential advantage of FIDA analysis compared to other interaction techniques is that it provides additional quality control information such as the R_h_ of the complex. Many designed proteins show undesirable multimerization, which should ideally be identified early in a screening pipeline. In this context, the experimentally determined R_h_ value can be compared to a predicted R_h_ from the PDB of the designed complex. All three of the positive binders in complex with FITC-ALFA were measured to a R_h_ within 0.2 nm compared to the predicted values (**Fig. 2E**).

### Screening directly on bacterial lysates by FIDA

Next, we aimed to test if it was possible to perform the initial binding screen on minimally processed bacterial lysates – thus potentially bypassing time-consuming protein purification steps. Various approaches are used for bacterial lysis in protein purification and direct lysate analysis. Among them, chemical and thermal lysis are some of the most efficient and least equipment-intensive, making them well suited for scalable applications.

First, we focused on chemical lysates, where bacteria are lysed by detergents followed by centrifugation to remove cell debris. Due to the high expression levels, the designed binders are the main protein components of such lysates (**Fig. 3A and Fig. S1**), although the purities and concentrations differ between variants. The concentration of proteins could in theory be approximated to allow normalization between binders prior to screening, yet we reasoned that adding steps to the selection pipeline defeats the overall purpose of saving time and resources by screening directly on lysates. However, it is still important to account for non-specific binding of lysate components, which can simply be addressed by inclusion of a preculture sample identical in composition but lacking the expressed binder.

**Fig. 3.**
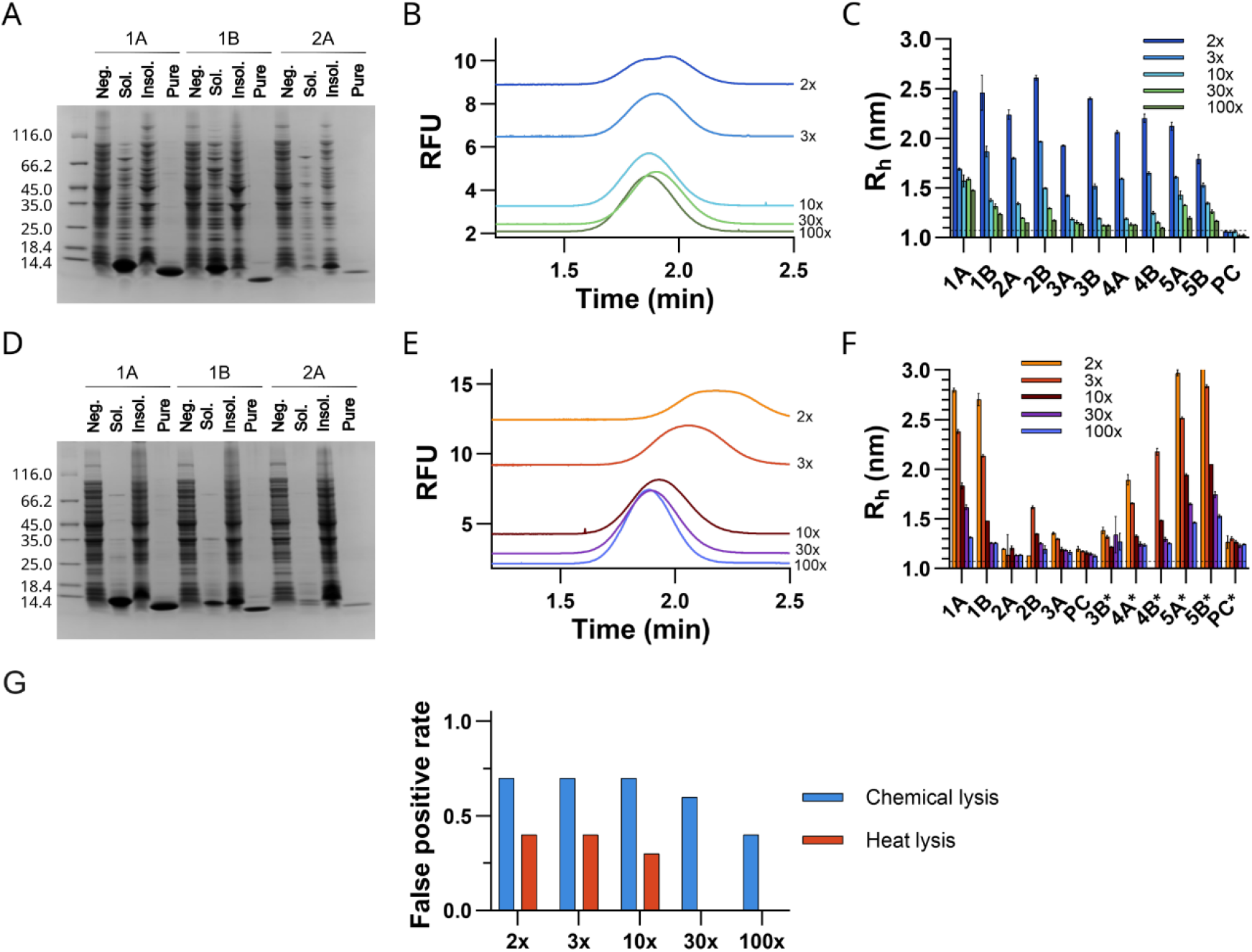
Binding screening in bacterial lysates. (A,D) Representative gel illustrating the purity of the samples used for FIDA screening (Sol.) in either chemical (A) or heat (D) lysates separated from insoluble material (Insol.) by centrifugation. A sample with no binder expression (Neg.) and one with purified binder (Pure) were loaded for comparison. (B,E) Examples of Taylorgrams obtained when using different dilutions of chemical (B) and heat (E) lysates, evidencing the high background fluorescence associated with the lower dilutions. In B, the Taylorgram for the 2x dilution additionally shows a double peak, which is a clear signal of buffer/baseline mismatch. A higher residence time is observed for the 2x and 3x diluted heat lysates (E) likely due to the increased viscosity of the samples. (C,F) Apparent size of FITC-ALFA (200 nM) when mixed with either chemical (C) or heat (F) lysates that were diluted to different extents. A sample of preculture (PC), i.e., lacking the expressed binder, was tested to account for non-specific binding to lysate components. The values correspond to mean ± SD (n = 3). The dotted lines mark the R_h_ of the indicator in clean buffer. The asterisks in F indicate that the samples were analysed in different days and should thus be compared to the corresponding preculture. (G) Rate of false positives in relation to the total number of binders for both lysis methods, where false positives correspond to binders that result in a 10% gain in R_h_ relative to the preculture.

Bacterial cell lysates exhibit high autofluorescence, which presents two challenges for FIDA-based screening. First, the increase in the background signal requires using a higher indicator concentration, to maintain an S/N > 50. Accordingly, 200 nM FITC-ALFA was used in the screening experiments, rather than the 50 nM used for purified binders. Second, the capillary mixing approach unavoidably introduces signal mismatch due to difference in the fluorescence baseline between the lysate and the indicator, the latter being in a clean buffer (**Fig. 3B**). If not accounted for, this will distort the shape of the Taylor peak and consequently the accuracy of the extracted R_h_. We identified three strategies to mitigate this problem: (i) increasing the indicator concentration even further to the point where this distortion is insignificant, (ii) pre-mixing the lysate and the indicator before analysis, and (iii) reducing the autofluorescence by diluting the lysates with a clean buffer solution. These strategies each have their own advantages and disadvantages, and the optimal choice depends on the specific application. In the following experiments, we chose to adopt the third option, as dilution offers the most effective approach in terms of time efficiency, sample consumption, and simplicity of setup. Notably, low dilution factors result in viscous samples which lead to an apparent increase in the calculated R_h_. Importantly, the apparent increase can be readily corrected by normalizing diffusivity based on viscosity - estimated from the difference in retention time relative to the indicator in buffer^15^.

To develop a screening protocol for interactions in the detergent-treated lysates, we measured the R_H_ of FITC-ALFA diluted into lysates serially diluted from 1:2 to 1:100 (**Fig. 3C**). In the least diluted lysates, the apparent hydrodynamic radii were bigger than that observed for the purified complex, suggesting that the indicator non-specifically interacts with a component of the lysate. Further dilution of the lysate with buffer led to a gradual decrease in apparent R_h_, the extent of decrease depending on the binder. Binder 1A was associated with a significantly higher R_h_ at the highest dilution factor, whereas binders 5A and 5B were approaching baseline levels, similar to the non-binders. As these interactions have similar affinities, the observed differences likely reflect varying expression levels of the binders (**Fig. 3A and Fig. S1**).

Instead of chemical lysis, the heat stability of the designed proteins offers an alternative strategy for preparing the bacterial lysates. In a heat lysis method, cell suspensions are heated to 95 ºC resulting in cell lysis, pelleting of most host proteins and inactivation of proteases. This method is often used for intrinsically disordered proteins^16^ but may also be a generally applicable strategy for thermostable, designed proteins. As we previously observed that the de. novo designed binders have a high thermal stability (**Fig. 2D**), we decided to use heat lysis to eliminate unstable proteins early in the workflow. Gratifyingly, heat lysis resulted in highly pure lysates for most of the binders tested, surpassing the purity achieved with the detergent-based lysis (**Fig. 3D and Fig. S2**). While heat lysis removes most protein contaminants, many cell metabolites are still preserved. Therefore, similar considerations apply to handling the auto-fluorescence as for the detergent-treated lysates, with increase in detection time particularly present in highly concentrated samples (**Fig. 3E**).

Similarly to the detergent-treated lysates, we screened a dilution series of the heat-treated lysates. The apparent R_h_ broadly follows the same pattern - higher than the expected complex size at the 1:2 dilution and gradually decreasing as the dilution factor increases **(Fig. 3F**). Noteworthy, the difference between binders and non-binders is greater in the heat-treated lysates than in the detergent-treated lysates at both high and intermediate concentrations. This suggests that there is less unspecific binding in the heat-treated lysates, likely due to the removal of most host proteins and the absence of detergent micelles.

In any project, the selection of a screening threshold will depend on the acceptable balance between false negatives and false positives. Using a 10% increase in R_h_ relative to the one measured for the preculture as the cutoff for a positive hit, all the previously validated binders were identified under both chemical and thermal lysis conditions. However, the false positive rate was highly dependent on the lysis method and dilution factor (**Fig. 3G**). While the false positive rate remained high across all dilutions for detergent-based lysis, it dropped to zero for heat lysis at dilutions of 1:30 and 1:100, providing a clear separation of binders and non-binders. Ultimately, the optimal threshold will depend on the R_h_ increase upon binding, the minimally acceptable affinity and the sensitivity to false positives and false negatives.

To further optimize the screening process, we developed a medium-throughput workflow for the expression and purification of de.novo designed protein binders using heat lysis (**Fig. S3**). In this method, the proteins are expressed in 2 mL bacterial cultures using 24-well plates, which are then subjected to heat lysis. Noteworthy, up to 96 bacterial cultures can be lysed in parallel by using a 96-well plate heating block. Centrifugation in a 96-well filter plate removes heat sensitive proteins, and thus most of the endogenous E¡.coli proteins, yielding highly pure lysates (**Fig. S3**) that can be directly used for screening with FIDA or further purified for next steps (e.g., affinity determination). A step-by-step protocol for this procedure is provided on protocols.io.

### FIDA for the evaluation of de.novo designed binders against complex targets

Next,.we aimed to assess whether FIDA is also useful for detecting binders to a larger target. For this purpose, we selected the guanylate kinase (GK) domain of the postsynaptic density protein 95 (PSD-95). The GK domain consists of three subdomains, the GMP-binding, the CORE, and the LID subdomains, with a total of 7 α-helices and 9 β-sheets^17^. Since the SH3 and GK domains are known to form a supramodule^18,19^, we chose to use a protein construct containing both domains for evaluating binding. Of note, assessing binding in this situation is inherently more challenging as the mean predicted change in R_h_ upon binding is much smaller (9-22%) than for the ALFA peptide (80-113%).

To design binders against the GK domain, we extracted its atomic coordinates from the crystal structure of PSD-95 GK in complex with a phospho-SAPAP peptide^20^ and used them as input for RFdiffusion. A set of hydrophobic residues (L552, Y580, Y604, Y609 and, for one of the design runs, L680) plus two arginine residues (R568 and R571) were selected as target residues. Binder designs were generated and scored as described for the ALFA-tag binders (**Fig. 2B**) resulting in a 7.7%.in.silico success rate prior to proteinMPNN resampling. The two sequences with lowest pAE_interaction_ from six diverse folds were selected for experimental testing (**Fig. 4A**).

**Fig. 4.**
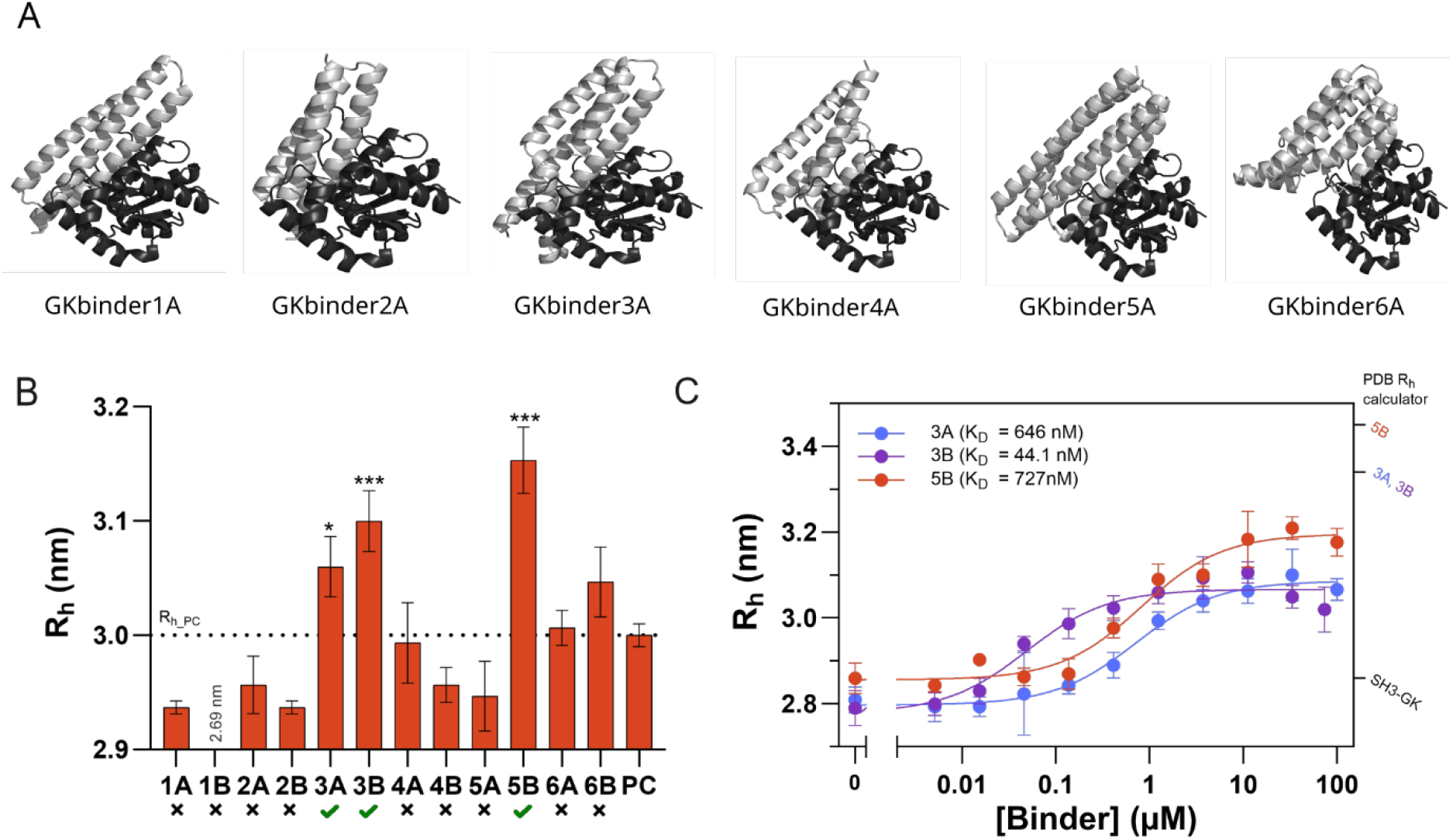
Screening and affinity determination of de. novo designed PSD-95 binders. (A) AlphaFold2 predicted structures of the PSD-95 GK domain (in black) in complex with the binders (in grey). For each binder backbone, there are two binder sequences (A and B), but only one of the complexes is shown here as they have a similar fold. (B) Binding screening in heat lysates from bacteria expressing the PSD-95 GK binders. 200 nM AF488-labelled SH3-GK was mixed with 30x diluted lysates inside the capillary and the resulting peaks used to determine its R_h_. A sample of preculture (PC) was tested to account for non-specific binding to lysate components. To compare the R_h_ obtained for the binders with the one of the PC (depicted by a dotted line), a one-way ANOVA Dunnett’s multiple comparisons test was performed (* statistically significant at p < 0.05; *** statistically significant at p < 0.001). Binders with a significantly higher R_h_ than the PC (green ticks) were considered as positive hits. (C) Titration curve of AF488-labelled SH3-GK (80 nM) with purified binders 3A, 3B and 5B. The solid lines show fitting to the 1:1 binding model. The predicted sizes for SH3-GK alone or in complex with the different binders are indicated in the right y-axis. All R_h_ values correspond to mean ± SD (n = 3)

All PSD-95 binders expressed in E¡.coli showed detectable protein expression and, as before, heat lysis yielded highly pure lysates (**Fig. S4**). Based on the results obtained for FITC-ALFA, we screened the binders at a 1:30 dilution using 200 nM AF488-labelled SH3-GK as indicator to ensure a S/N > 50. This resulted in a R_h_ of 2.96 nm, closely matching the one predicted for the structure of SH3-GK without AF488 (2.86 nm). Since the expected size change for SH3-GK upon interaction with any of the designed binders is significantly smaller than for ALFA-tag - with the minimum predicted R_h_ change being 7.9% - we considered all binders that induced a statistically significant increase in R_h_ compared to the negative control as positive hits (**Fig. 4B**). Binders 3A, 3B and 5B met this requirement.

To confirm whether binders 3A, 3B and 5B were true hits and obtain their binding affinity towards the SH3-GK supramodule, the binders were purified and titrated against AF488 labelled SH3-GK (**Fig. 4C**). Fitting the curves to a 1:1 binding model resulted in K_D_ values of 646 ± 108 nM, 44.1 ± 16 nM and 727 ± 216 nM for 3A, 3B and 5B, respectively, confirming that they bind to the target protein. Comparison to the R_h_ values calculated from the AlphaFold3 predicted complex structure suggested that the complexes have the expected sizes. As we aimed to validate the screening on bacterial lysates as a strategy for initial hit identification, we purified the remaining binders and performed a titration experiment in the same manner (**Fig. S5**). The results revealed that one binder with low micromolar affinity, binder 5A, was missed - likely due to its very low expression level (**Fig. S4**). Nonetheless, these findings support the use of bacterial heat lysates for initial binding screening, even when the target exhibits only a modest increase in R_h_ upon binding.

## Discussion

In this work, we test the potential of FIDA as a tool for screening computationally designed protein binders. Using two protein targets, differing in both size and structure, we have shown that FIDA can detect binding in a single measurement directly in bacterial lysates with minimal sample pretreatment.

Practical implementation of a FIDA-based binder screening pipeline involves compromises between sensitivity and throughput and requires balancing trade-offs between false positives and false negatives. We aimed to establish the fastest possible protocol that identifies promising designs worth further investigation. We found that the most effective strategy is to screen for binding using a single capmix experiment with a 30-fold dilution of heat-treated lysate. This excludes all false positives without sacrificing the detection of true positive hits. However, this approach has some drawbacks that make it unsuitable for certain applications. First, it inherently selects for both binding and heat-stability. This works well for the small helical bundles but may exclude more complex binder designs that are not thermostable. In such cases, screening in detergent-treated lysates is still a feasible alternative at the expense of purifying and titrating a higher number of binders. Second, the absence of normalization to protein expression-levels means that high affinity binders with poor expression might be missed. A pragmatic approach to handling this is to assess the expression-levels of binders near the detection threshold and only include those with low expression for follow-up analysis. Nevertheless, one could argue that this “pick-the-winners” approach with single-point lysate screening allows rapid and efficient identification of binders with both high affinity, robust expression and thermostability.

Designs cannot be neatly categorized into binary categories of binders and non-binders. For each application a subjective choice of what is a “good enough” affinity should be made, which affects the appropriate threshold. We see indications of weak binding in some of the discarded binders, however in each case the retained proteins have higher affinity suggesting the threshold works in ranking binders. The filtering involves two subjective parameters that affect the stringency of the screening: the dilution factor and the R_H_ threshold. We recommend that these parameters should be optimised on a case-by-case basis, depending on the properties of the binders and the target. Initially, if too many hits are identified in a pilot experiment, the dilution factor could be increased to identify the strongest hits. Secondly, the R_h_ threshold should be adjusted for each target based on the expected change in R_h_ upon binding. In the case of the PSD-95 binders, a fixed R_h_ threshold of 10% would result in missing even a high-affinity binder with a K_D_ of 44 nM. Instead, we suggest using a statistical test, where binders are considered hits if they produce a significant increase in R_h_ relative to the indicator in blank lysate.

FIDA has some advantages compared to BLI, one of the main methods for binder screening. First, FIDA does not require immobilization of the target protein. Three important advantages arise from this: (i) non-specific interactions with the sensor surface are avoided, (ii) no steric effects are introduced that could hinder binding to the protein target and (iii) optimization is kept to a minimum. Furthermore, FIDA uses size measurements as a quality control parameter, providing information not only on binding, but also information about the structure, stoichiometry and heterogeneity of the sample.

The most important limitation of screening using FIDA is its reliance on size change to assess binding. This works well when the binder is approximately the same size as the target. However, it limits its sensitivity when the difference in R_h_ between the target and the complex is not significant, which typically is the case for larger target proteins. The expected change in R_h_ should therefore be estimated in advance, to determine if FIDA is an appropriate screening method.

In conclusion, FIDA stands out as a straightforward technique for screening de.novo designed binders, allowing the identification of positive hits within one day of protein expression and with low material consumption. Its strengths and disadvantages are orthogonal to biosensor techniques as BLI and thus represent a useful addition to protein design screening efforts.

## Acknowledgements

This work was supported by the Danish National Research Foundation (DNRF133) awarded to the Center for Proteins in Memory (PROMEMO) and the Independent Research Fund Denmark (026-00069B) to MK, and a postdoctoral fellowship from the Lundbeck Foundation to FP (R449-2023-1396), and an instrument grant from the Carlsberg Foundation to Jørgen Kjems (CF20-0610).

## Author contributions

FP, JSN and MK conceived and designed the study. FP, JSN and EZ performed the FIDA experiments. FP, EZ and ECP purified the proteins. IG and VL optimized the protocol for mid-throughput expression and purification of de.novo designed binders. MK designed the binders. FP, JSN and MK supervised the experiments. FP, JSN and MK wrote the original draft with contributions from ECP and IG. All authors have given approval to the final version of the manuscript.

## Notes

### Competing Interest Statement

The authors have declared no competing interest.

