## Supplemental information for "Screening de. novo designed protein binders in unpurified lysate using flow induced dispersion analysis"

**Protein sequences**

**Color coding:**

6xHis-tag

Thrombin or TEV cleavage site

Binder

**ALFA-tag binder 1A**

MGSSHHHHHHSSGLVPRGSHMHMSEEFRKEIEERMERIRERLLRALETADLETLLALALEEVEELKEYYESLSEEEQEILFERLDEFFARLRAELAALTPELRAKALAALAALAEELA

**ALFA-tag binder 1B**

MGSSHHHHHHSSGLVPRGSHMHMSAAFREALRARAEARLEAFLEALRTKSVEELLALARAEVAELKELYASLSEEEQEILLEILDEAAERLRARLAELSPELRAKALEVLARLAEELA

**ALFA-tag binder 2A**

MGSSHHHHHHSSGLVPRGSHMHMSRREEIRARIEAELAAILRRIGARMESPELEPAIIALAARGYRELLALVEEDPELAERTRDARLAELEAELLALV

**ALFA-tag binder 2B**

MGSSHHHHHHSSGLVPRGSHMHMSRRDRIRAEVLARVAALLRRIGAEMADPALEPAVIRRAAAGLEELFALIDADPDAAAATLDARLAALEAELRALV

**ALFA-tag binder 3A**

MGSSHHHHHHSSGLVPRGSHMHMVTSPLELFLLLEEAFRELAEEEGDPERQERRRERLLELALELLGDDPRVERAARAARLLVDAERAGAPEEELAALRAALLAELA

**ALFA-tag binder 3B**

MGSSHHHHHHSSGLVPRGSHMHMVTSPLELYLLLYEAFRELAEEIGDEEEFEREAERLLALALEILGDDERVRLAYERARALVEAERAGLPAEELAALRAAVLAELA

**ALFA-tag binder 4A**

MGSSHHHHHHSSGLVPRGSHMHMSALERLRAAIEAEIKWKEEQAKKAEELGDKESAELFREIAELLETLYEFVELALEGEVERARRGFELALELALEAEREAAERLLEKFGNKEEAEEIRARAEASAEELRALFAELLA

**ALFA-tag binder 4B**

MGSSHHHHHHSSGLVPRGSHMHMSALEELKELLEKHIEWLKEKAKESKELGDEESAELFEEMAELTKTLFEFVELIEEGEVERGRRGFELALELLREAAKEAAELHKKKFGDEERAKEIIAYADDFTKRLTEKVEELIA

**ALFA-tag binder 5A**

MGSSHHHHHHSSGLVPRGSHMHMAEEELERLKEEILERVREVLESLSPEEREALERLSPEEALDRVLERMAELDEESRELAERARELLE

**ALFA-tag binder 5B**

MGSSHHHHHHSSGLVPRGSHMHMEEEERRRLEEEILEAVREALAALPPEERAALEELSPEEALDRVLELMAERDERSRALAEAARRLLE

**PSD-95 GK-domain binder 1A**

MGSSHHHHHHSSGLVPRGSHMSEVEEASLELDELIDQMERLLERAEEALEEAEKALKEGDLETAEKLLKVVEDHLRRFNSLYTRIGNELKKLPPELQAEYQKRIDELEERRLEILKRLAEL

**PSD-95 GK-domain binder 1B**

MGSSHHHHHHSSGLVPRGSHMEEVEEATEELNELLDQLERLLEIAEKNLEEAKKLLEEGDLESAERKLKIVEDHIRRFYSLYTRIGNELKKLPPEQQKIFNEKIDKLEERRLKILKELAEL

**PSD-95 GK-domain binder 2A**

MGSSHHHHHHSSGLVPRGSHMSAEKEAFLNGARVLSSALRKMLRERLKEYKETGSEEAAKYAEEVLKEMRELAEDLEKLGFEVEAVELKERIEEYEKELKKLKE

**PSD-95 GK-domain binder 2B**

MGSSHHHHHHSSGLVPRGSHMSAEKEAFYNGARVLWSALRKMARERLKEYKETGSKEAAEYLKRVLKEMRELAEDLRKLGFDVEATELEERIKEYEKELKELEK

**PSD-95 GK-domain binder 3A**

MGSSHHHHHHSSGLVPRGSHMFEARLQAALAAIGELADLYEDLYLETRKAYAEIKASKDWKERLKIVKELLKYVKEKMEAGLAREDEILVDAEAVLQEAYDRAPEEAESLESPASQIRGALLIKSSARHNIIELLHELKKLFAKDAEKPEAQEGKKLIEEIEKLLKGP

**PSD-95 GK-domain binder 3B**

MGSSHHHHHHSSGLVPRGSHMMQEELEAAKAAIGKLADTAEDLYLRTLEAYEKIKASTDWEEKLAITQELLAYVQAVMAAGYAERDAVLVDAEAVIQKAEEAAPEEAKSLESLTSEIKGALLIASSARSRIIELLHEMKKLFAKDAEHPAAQQGLELIDQILATLKGP

**PSD-95 GK-domain binder 4A**

MGSSHHHHHHSSGLVPRGSHMEKERREELFERYVRRAREIIERYKKIKPELSDDSEELRELVLEALDEVARAAILAGLKIYPLLIQRMLLVEKGDVLDEEALDYAERALERIEKKREEQRKEEE

**PSD-95 GK-domain binder 4B**

MGSSHHHHHHSSGLVPRGSHMAEAARRAALEAALAAARATLARLEALKPLLSDDSQELRDLAGDALDHAARAAILAGLDIHPLLVQRMLLLEKGDILVEEAVDYVRRVLERIEAKRAEQEAAAA

**PSD-95 GK-domain binder 5A**

MGSSHHHHHHSSGLVPRGSHMMIEEVEKKLKEMLEFSTKFMKELAKKFLEHLKKLVEKYGTEEIKKRAEKLKEFLDEFLNDFEDFLKRMFDAIIKYFKEGNIEKAAKVALDLARLFFRRAVRRVEEVKKEIEISTGFEGLVDAVVDYVMLVANTFAKNVDLEPLEIVEKMMEKLKEAAEKAEKELEKAMKKEEEEIEKKL

**PSD-95 GK-domain binder 5B**

MGSSHHHHHHSSGLVPRGSHMDIEEVKEKLKELLEKSLAFFKELADKVRAGLAAIVAETGTEEVARNAAILNEAIDELLNDIEDHLKRTIDAIIKYIEEGDIETAVKVALDLSRLFSRRAVRKVEEVRKESPLSTGFEITVDAIVDHIMRVANTFYKNKDLPPLERIEKVLEVLDESAKEAAARTDEGLAAYRAEVAARL

**PSD-95 GK-domain binder 6A**

MGSSHHHHHHSSGLVPRGSHMMEREEDLLLRRLREEGKEEEAEELELKLALELARSVLDLVEALAKALAKGDLQSAADIARLKMMSSDKTVHDLAEIFLRYMSALLEDPAAAGEVFTTDLEALLARYAPNPEVSALLQALLDAYRAAAGGPLATLAAALAAYAKWLEERAKELEKKLKE

**PSD-95 GK-domain binder 6B**

MGSSHHHHHHSSGLVPRGSHMKMRERDRLLRELREAGRLEEARRLELEEALELAEAVLPLVEALAKALEKGDYESALQIARLKMMSSNKTVHDIAEIFMRYMSALLQDPAKAAEVFTTALTELREKHAPNPEVAAILDALLEAYKAAAGKSPAELAAALRAYAEWLAARAAEARAELAA

**PSD-95 SH3-GK module**

MGSSHHHHHHSSGLVPRGSHMENLYFQSGSFYIRALFDYDKTKDCGFLSQALSFRFGDVLHVIDAGDEEWWQARRVHSDSETDDIGFIPSKRRVERREWSRLKAKDWGSSSGSQGREDSVLSYETVTQMEVHYARPIIILGPTKDRANDDLLSEFPDKFGSCVPHTTRPKREYEIDGRDYHFVSSREKMEKDIQAHKFIEAGQYNSHLYGTSVQSVREVAEQGKHCILDVSANAVRRLQAAHLHPIAIFIRPRSLENVLEINKRITEEQARKAFDRATKLEQEFTECFSAIVEGDSFEEIYHKVKRVIEDLSGPYIWVPARERL

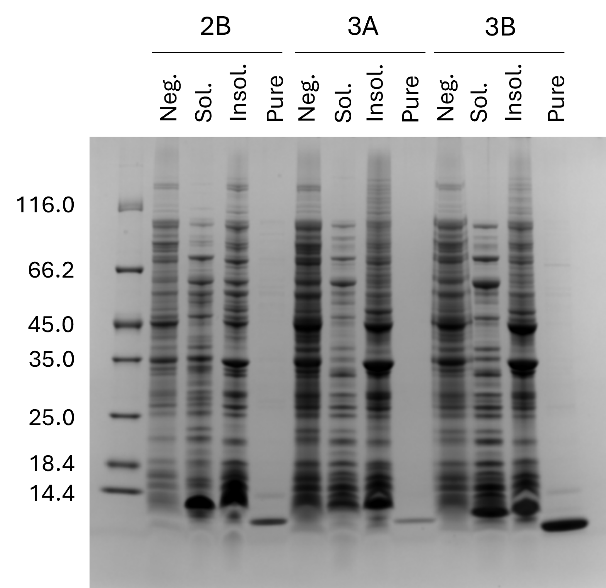

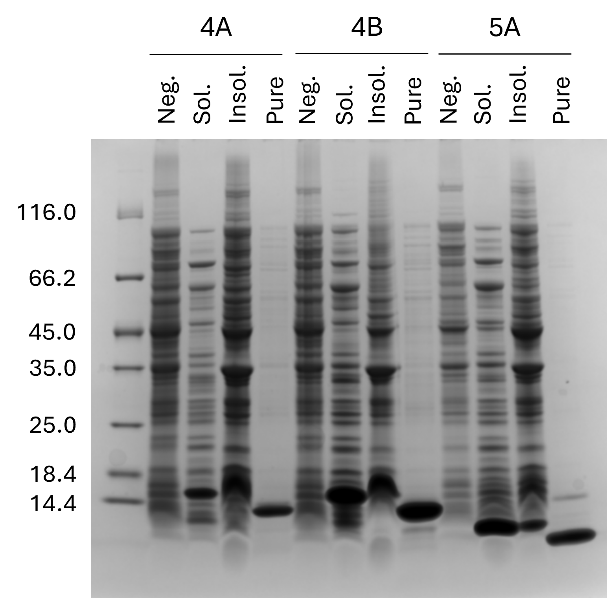

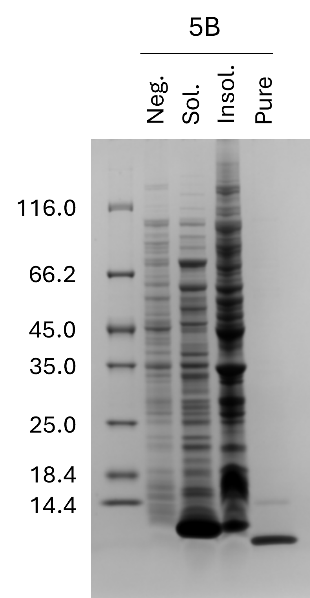

**Fig. S1. SDS-PAGE analysis of soluble and insoluble fractions after chemical lysis for ALFA-tag binders*.*** *E.coli* cells expressing binders 2B-5B were lysed with BPER reagent and the proteins present in the soluble (Sol.) and insoluble (Insol.) fractions visualized by SDS-PAGE. Cells with no expression of the protein (Neg.) and a sample of purified protein (Pure) were loaded for comparison. The soluble fractions were used for FIDA measurements.

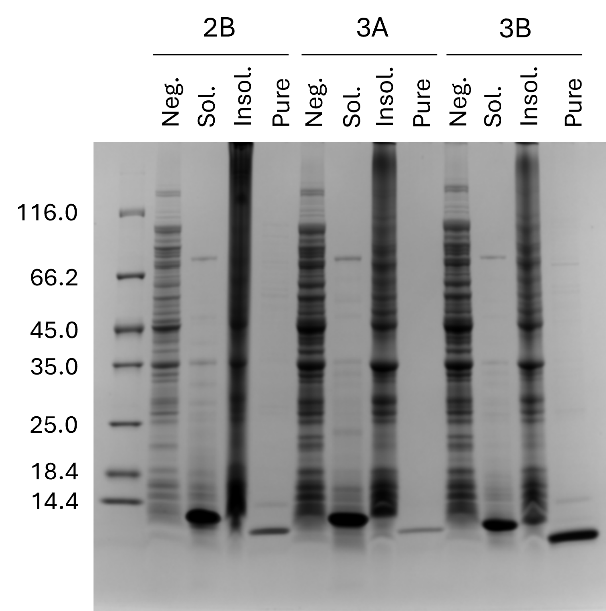

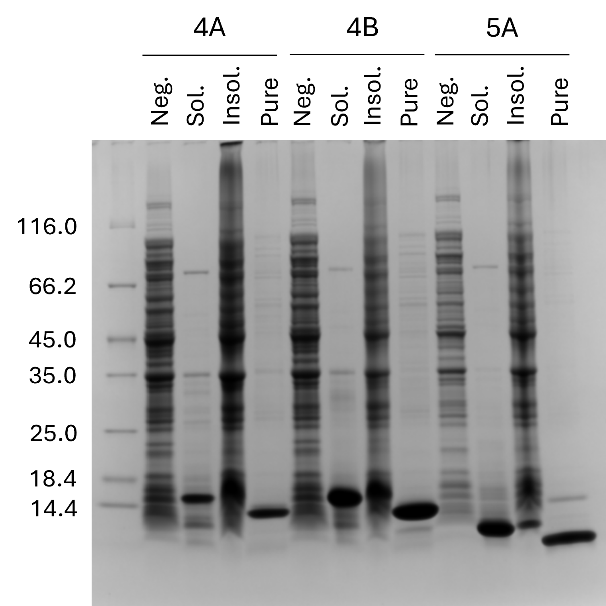

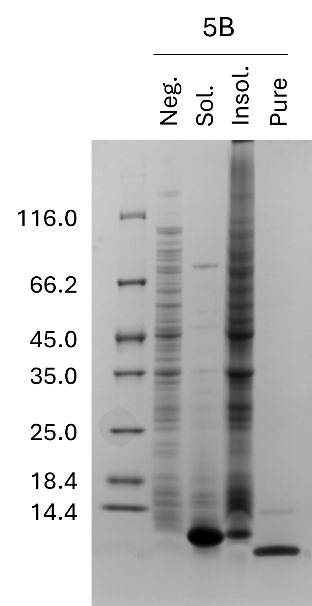

**Fig. S2. SDS-PAGE analysis of soluble and insoluble fractions after heat lysis for ALFA-tag binders*.*** *E.coli* cells expressing binders 2B-5B were subjected to heat lysis and the proteins present in the soluble (Sol.) and insoluble (Insol.) fractions visualized by SDS-PAGE. Cells with no expression of the protein (Neg.) and a sample of purified protein (Pure) were loaded for comparison. The soluble fractions were used for FIDA measurements.

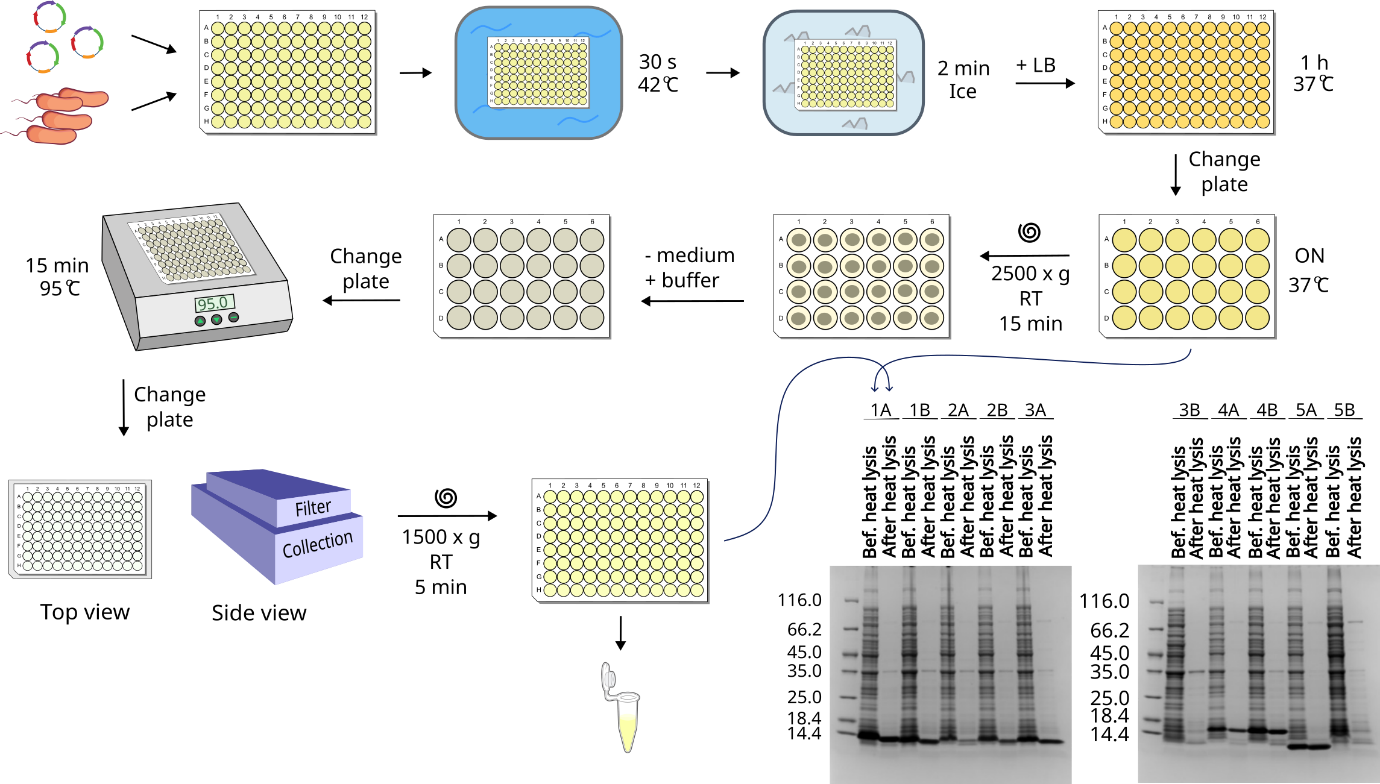

**Fig. S3. Overview of the protocol for expression and purification of *de novo* designed proteins using heat lysis.** ALFA-tag binders were expressed and the cultures processed following this protocol. Samples before and after heat lysis were taken and loaded on SDS-PAGE gels to compare their purity. A detailed protocol for this procedure can be found in protocols.io.

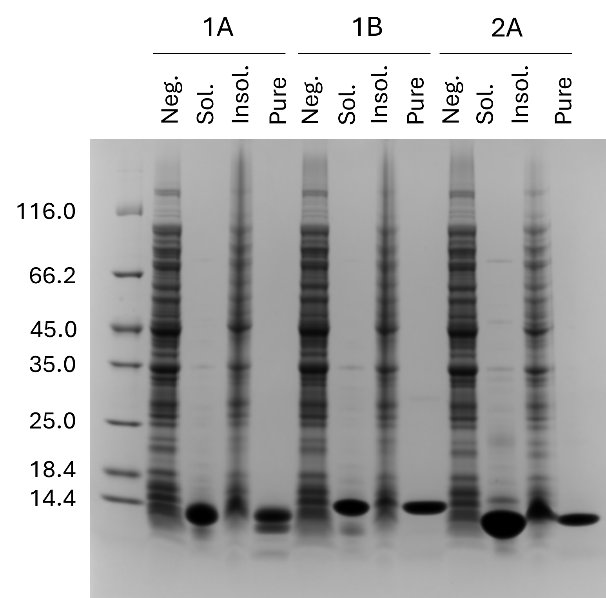

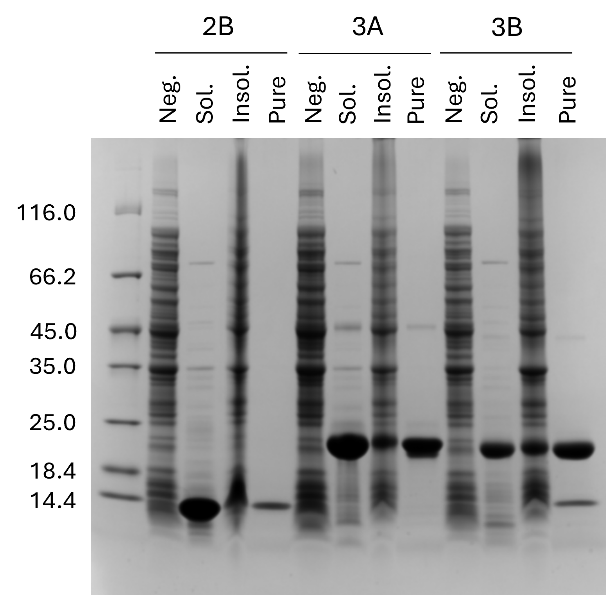

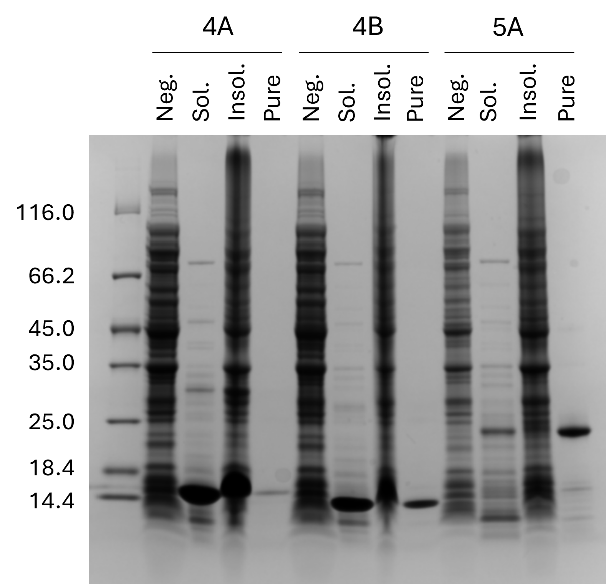

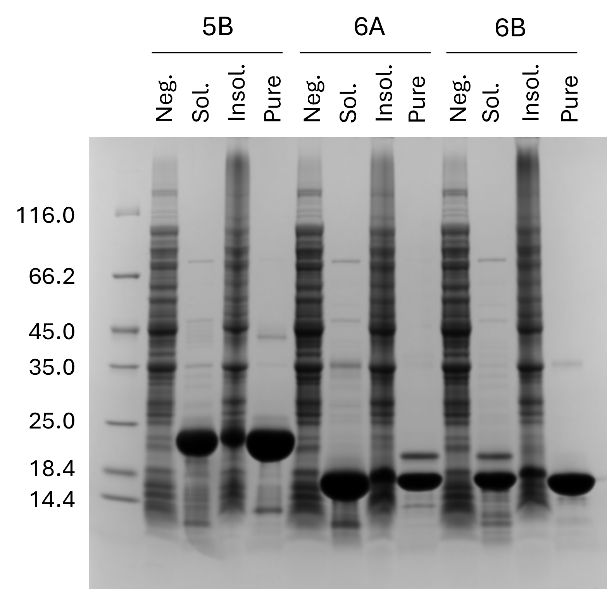

**Fig. S4. SDS-PAGE analysis of soluble and insoluble fractions after heat lysis for PSD95-GK domain binders*.*** *E.coli* cells expressing binders 1A-6B were subjected to heat lysis and the proteins present in the soluble (Sol.) and insoluble (Insol.) fractions visualized by SDS-PAGE. Cells with no expression of the protein (Neg.) and a sample of purified protein (Pure) were loaded for comparison. The soluble fractions were used for FIDA measurements.

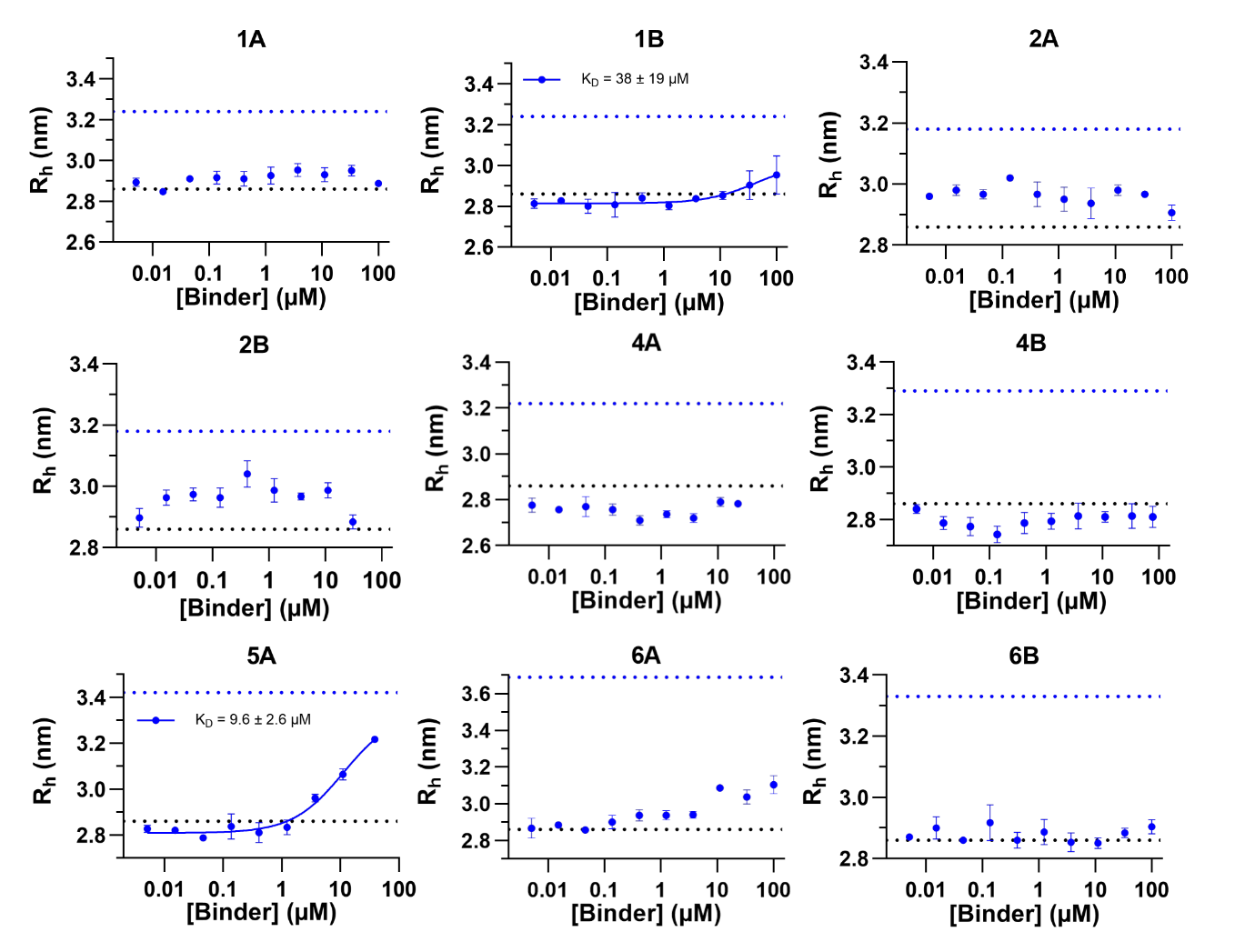

**Fig. S5. Titration curves of AF488-labelled SH3-GK with purified GK-domain binders.** Apparent R_h_ of AF488-labelled SH3-GK (80 nM) as a function of binder concentration. The R_h_ values represent mean ± SD (n = 3). In the case of binder 1B and 5A, the values could be fitted to a 1:1 binding isotherm, which is represented as a solid blue curve, allowing to calculate the K_D_. The black and blue dotted lines illustrate the predicted size of the SH3-GK domain alone and in complex with the different binders, respectively.
